# CTP-Synthase 2 from *Arabidopsis thaliana* is required for complete embryo development

**DOI:** 10.1101/2021.01.06.425551

**Authors:** Daniel Hickl, David Scheuring, Torsten Möhlmann

## Abstract

Pyrimidine de *novo* synthesis is an essential pathway in all organisms. The final and rate limiting step in the synthesis of the nucleotide CTP is catalyzed by CTP-Synthase (CTPS) and Arabidopsis harbors five isoforms. Single knockouts of each of these do not show apparent phenotypical alterations with the exception of *CTPS2.* T-DNA insertion lines for this isoform were unable to produce homozygous offspring. Here we show that *CTPS2* exhibits a distinct expression pattern throughout embryo development and loss of function mutants were embryo lethal, as siliques from *+/ctps2* plants contained nearly 25 % aborted seeds. This phenotype was rescued by complementation with *CTPS2* under control of its endogenous promoter. Reporter lines revealed *CTPS2* expression only in the tip of columella cells in embryos of the heart and later stages. Furthermore *CTPS2* expression in roots, most pronounced in the columella cells, shoots and vasculature tissue of young seedlings was observed. Filial generations of *+/ctps2* plants did not germinate properly, even under external cytidine supply. During embryo development *CTPS2* expression was similar to the well known auxin reporter DR5. Indeed, the cloned promoter region we used in this study possesses a repeat of an auxin response element. Thus, we conclude that CTPS2 is essential for CTP supply in the developing embryo and a knockout of *CTPS2* is embryo lethal.

## Introduction

Nucleotides are essential building blocks for the production of nucleic acids. In addition nucleotides represent the main energy carriers in biochemical reactions, function as nitrogen, carbon or phosphate source under nitrogen limiting conditions and as co-factors in phospholipid biosynthesis. Due to their chemical structure nucleotides are divided in purines and pyrimidines (Moffatt and Ashihara 2002; Moore 1993; Zrenner et al., 2006). In plants, nucleotide metabolism consists of i) *de novo* synthesis, ii) salvage and iii) degradation (Zrenner et al., 2006). Pyrimidine *de novo* synthesis consists of 6 enzymatic steps distributed to chloroplast, cytosol and mitochondria, ending up with the production of Uridine monophosphate (UMP) in the cytosol. This intermediate is phosphorylated by UMP kinase to UDP. Uridine mono-, di- and triphosphates are equilibrated by nucleoside diphosphate kinases. The last step of pyrimidine *de novo* synthesis is the amination of UTP to CTP, conducted exclusively by CTP-synthases (CTPS) (Zrenner et al., 2006; Witz et al., 2012; Moffatt und Ashihara 2002). CTP-synthases represent a conserved enzyme family found across kingdoms. The demand of CTP as part of DNA is especially high during cell division and in developing tissues. Therefore CTPS activity was described to be regulated on different levels, e.g. posttranslationally (phosphorylation), allosterically (GTP) or feedback inhibited by its product (CTP) (Levitzki and Koshland 1972; Chang and Carman 2008). A further enzymatic regulation is the polymerization to filaments, which was studied in several organisms like *E. coli, S. cerevisiae. D. Melanogaster* and *H. sapiens* (Barry et al., 2014; Liu 2010; Lynch et al., 2017; Noree et al., 2014). Plant CTPS were first described in *Arabidopsis thaliana* (hereafter referred to as Arabidopsis) by Daumann et al., 2018, describing features of five isoforms including the ability of filament formation. The isoforms show tissue specific expression patterns, which are dynamic for CTPS1 and 4 under abiotic stresses (Daumann et al., 2018; Hruz et al., 2008). However, single knockout mutants did not show any phenotype under long or short day regime, except CTPS2. We were not able to produce homozygous T-DNA insertion lines for this isoform, indicating a special function during embryo development or germination (Daumann et al., 2018).

Since the 6 core enzymatic steps of pyrimidine *de novo* synthesis are facilitated by 5 enzymes, each encoded by one gene, homozygous knockout lines cannot be generated in this pathway (Schröder et al., 2005). Antisense lines of aspartate transcarbamoylase (ATCase) and dihydroorotase, which facilitate the second and third step in pyrimidine *de novo* synthesis, cause a smaller phenotype in potato. Nevertheless, these antisense lines show a smaller, but viable phenotype, which is even able to produce tubers when the protein level is 20 % of the corresponding wildtype. Furthermore, no changes in nucleotide pools were observed in fully developed tissues unless expression dropped below a threshold, pointing to an efficient maintenance by recycling pathways in older plants (Schröder et al., 2005). Additionally to the knockout of genes in the pyrimidine *de novo* synthesis, knockout of genes encoding for enzymes in the plastidial pentose phosphate pathway, which products and intermediates are necessary for purine *de novo* synthesis hampers nucleotide production. Inhibition of purine and pyrimidine synthesis by knockout is embryo lethal, underlining the importance of nucleotide homeostasis in developing embryos (Andriotis and Smith 2019). Thus a large set of genes encoding for enzymes in the nucleic acid production pathway can be classified as essential for embryo development *(EMB* genes) (Meinke 1985). The analysis of transcriptional regulation by TOR mediated growth signals identified ATCase, DHODH and CTPS as target in nucleotide synthesis, underlining their relevance in regulating central metabolism (Xiong et al., 2013, Busche et al., 2020). Due to the fact, that pyrimidine *de novo* synthesis is facilitated by enzymes, which are mainly encoded by one gene it is not surprising, that homozygous knockout of any of these genes is embryo lethal. Since *A. thaliana* harbors five CTP-synthases it was surprising, that no homozygous knockout for isoform 2 could be generated (Daumann et al., 2018).

Here we report, that CTPS2 is indispensable for proper embryo development. Although CTPS2 expression is strongly controlled in roots of seedlings, the production of CTP by CTPS2 is critical during embryo development, as we showed by the number of aborted seeds before and after complementation, as well as the tissue specific expression in developing embryonic cells.

## Material and Methods

### Plants and growth conditions

The analysis used Arabidopsis thaliana variety Col-0 as WT and *CTPS2* (At3g12670) T-DNA insertion lines from the GABI-Kat collection (Kleinboelting et al., 2012). These were GABI_032C02 and GABI_156G07, designated as *+/ctps2-1* and *+/ctps2-2,* respectively. Seeds of soil grown plants were sowed on standardized ED73 soil (Einheitserde und Humuswerke Patzer, Buchenberg, Germany), incubated for 48 h at 4 °C for stratification and transferred to growth chambers under long day regime (14 h light/ 10 h dark). Growth conditions were: light intensity 120 μmol quanta m-2 s-1, temperature 22 °C, humidity 60 %. Segregation and promoter analysis were conducted with surface sterilized seeds on ½ MS agar plates (Murashige and Skoog, 1962), supplemented with vitamins, 1 % sucrose (w/v), 0.05 % MES-KOH (w/v), pH 5.7 and 0.8 % agar. Seeds were surface sterilized by addition of 500 μl 70 % Ethanol supplemented with 0.1 % Triton X-100 for 5 min on an end-over-end shaker. After 1 min centrifugation at 5000 g for 1 min, supernatant was discarded and seeds were washed twice with 100 % Ethanol for 1 min. The seed/ethanol solution was immediately pipetted on a sterile filter paper and dried in the airflow of a sterile bench.

### Construction of CTPS2 reporter and complementation lines

The CTPS2 promoter activity analysis used 1002 bp upstream from exon 1, while the complementation of *+/ctps2* plants used the full promoter (1864 bp) and protein coding sequence of *CTPS2* as given on the Aramemnon homepage (Schwacke et al., 2003). Primers used are given in Table S1. WT gDNA was used as template for the amplification of the *CTPS2* promoter sequence. Subsequently to PCR the amplified sequence was separated on an agarose gel (1 % w/v) and isolated by gel digestion with a Nucleo-Spin^®^ Extract II kit (Macherey Nagel, Düren). Afterwards att-sites were attached by another PCR, the amplificate isolated as described above and used for Gateway cloning. Once integrated in the entry vector pDONR (Thermo Fisher Scientific, Waltham Massachusetts USA), pBGWFS7.0 was used as target vector for *CTPS2* promoter activity analysis and pMDC123 for complementation of *+/ctps2* plants with the *CTPS2* full length construct (Curtis and Grossniklaus, 2003; Karimi et al., 2002). Correct insertion of the cloning procedure was checked by enzyme digestion and Sanger sequencing (Seq-IT, Kaiserslautern Germany).

*CTPS2* reporter construct was transformed into WT plants and *CTPS2* full length sequence into *+/ctps2* plants according to the floral inoculation procedure (Narusaka et al., 2012).

### Primers used

This study used primers for the amplification of *CTPS2* promoter and full length sequence as well as Gateway cloning of these sequences. Furthermore, T-DNA insertion was analyzed according to recommended primers by the GABI-Kat Homepage (Kleinboelting et al., 2012). During T-DNA insertion analysis, primers for EF1α were used as gDNA quality control. Sequencing was conducted with primers used for amplification of the 1002 bp of the *CTPS2* promoter and for the complementation construct also with sense primers from position 700 to 5600 (Table S1).

### Light microscopy and sample preparation

Siliques of soil grown plants (WT and *+/ctps2-1)* were harvested 8-10 DAF, longitudinal dissected and placed on a microscope slide, covered with parafilm. Seed shape was investigated on 12-15 DAF siliques, which were dried for up to 7 days at room temperature. Siliques were carefully opened under the light microscope and placed on a microscope slide covered with parafilm for better grip. Herein WT, *+/ctps2* plants and *+/ctps2* plants complemented with endogenous CTPS2 were used. Images were taken with a Leica MZ10 F microscope equipped with a Leica DFC420 C camera.

### Confocal laser scanning microscopy and sample preparation

Developing siliques of CTPS2::GFP promoter lines were harvested 1.5-8 DAF, dissected, seeds isolated and transferred to 1.5 ml reaction tubes. After addition of 10 μl ddH2O per seed, embryos were carefully squeezed out with a potter. The embryo/water solution was transferred to microscope slides and immediately used for microscopy. For the investigation of roots of CTPS2::GFP plants, seeds were placed on ½ MS agar plates containing 1 % agar and vertically positioned in the growth chamber. At the indicated time points seedlings were mounted in propidium iodide (PI) solution (0.01 mg/ml) for cell wall staining, followed by confocal laser scanning microscopy. Images in Figure 3, 5 and 6 were taken with a Leica TCS SP5II microscope. The excitation for GFP was 488 nm and detection of emission was at 505-540 nm. Chlorophyll autofluorescence of embryos and PI fluorescence was detected at 514 nm emission and at 651-704 nm excitation through a HCX PL APO 63 x 1.2 W water immersion objective.

Images in figure 4, S1 and S4 were acquired with a Zeiss LSM880, AxioObserver SP7confocal laser-scanning microscope (INST 248/254-1), equipped with a Zeiss C-Apochromat 40×1.2 W AutoCorr M27water-immersion objective with fluorescence settings as given in (Kaiser et al., 2019).

Images were processed using Leica software LAS X (version 3.3) or Zeiss software ZEN 2.3.

### Auxin treatment

Surface sterilized seeds of CTPS2::GFP reporter plants were placed on ½ MS agar plates and vertically grown for 7 days in a long day chamber. Afterwards seedlings were transferred for 20 h to ½ MS agar plates containing 250 nM NAA. NAA was solubilized in 100 % DMSO and add to lukewarm ½ MS media prior to pouring the plates. As control 250 nM DMSO was added to another batch of plates and the same number of seedlings was transferred to control conditions as for NAA treatment.

Fluorescence was detected with a Zeiss LSM880 as mentioned above. Images were processed using ImageJ (version 1.51j8) and the mean grey value is given from fluorescence of 10 trichoblast cells of 5 biological replicates.

### Statistical analysis

All experiments were carried out at least three times. Box limits in the graphs represent 25th to 75th percentile, the horizontal line the median, small light grey box the mean and whiskers minimum to maximum values.

## Results

### Cytidine supplemented Segregation analysis of *+/ctps2* seeds

Previously, we identified two *CTPS2* T-DNA insertion lines from the Gabi-Kat collection, which were unable to produce homozygous offspring (Daumann et al., 2018). These heterozygous lines GK_032C02 and GK_156G07 were designated as *+/ctps2-1* and *+/ctps2-2,* respectively. A former Segregation analysis of these two lines showed a deviant germination pattern, compared to the expected 1:2:1 ratio for WT:heterozygous:homozygous offspring (Daumann et al., 2018). Therefore, we concluded, that the crucial function of CTPS2 is probably in germination and/or embryo development. To confirm the before mentioned irregular pattern, we repeated the experiment in our study and found that nearly 10 % of *+/ctps2-1* and *+/ctps2-2* seeds, which were sown on ½ MS agar plates did not germinate compared to WT (Figure 1A and C), implicating a homozygous *CTPS2* knockout in this seeds. Since nucleotide *de novo* synthesis is very energy intensive, plants are able to recycle nucleosides via the salvage pathway (Zrenner et al., 2006). On this occasion, external supplied nucleosides can be imported by the equlibrative nucleoside transporter 3 (ENT3) for being phosphorylated to nucleotides (Traub et al., 2007). The supplementation of ½ MS agar plates with 1 mM cytidine for another segregation analysis results again in 10 % non-germinated seeds of both *+/ctps2* lines, whereas WT seeds grew normally (Figure 1B and C). This finding suggests that the critical role of CTPS2 takes place in seed production rather than germination.

**Figure 1.**
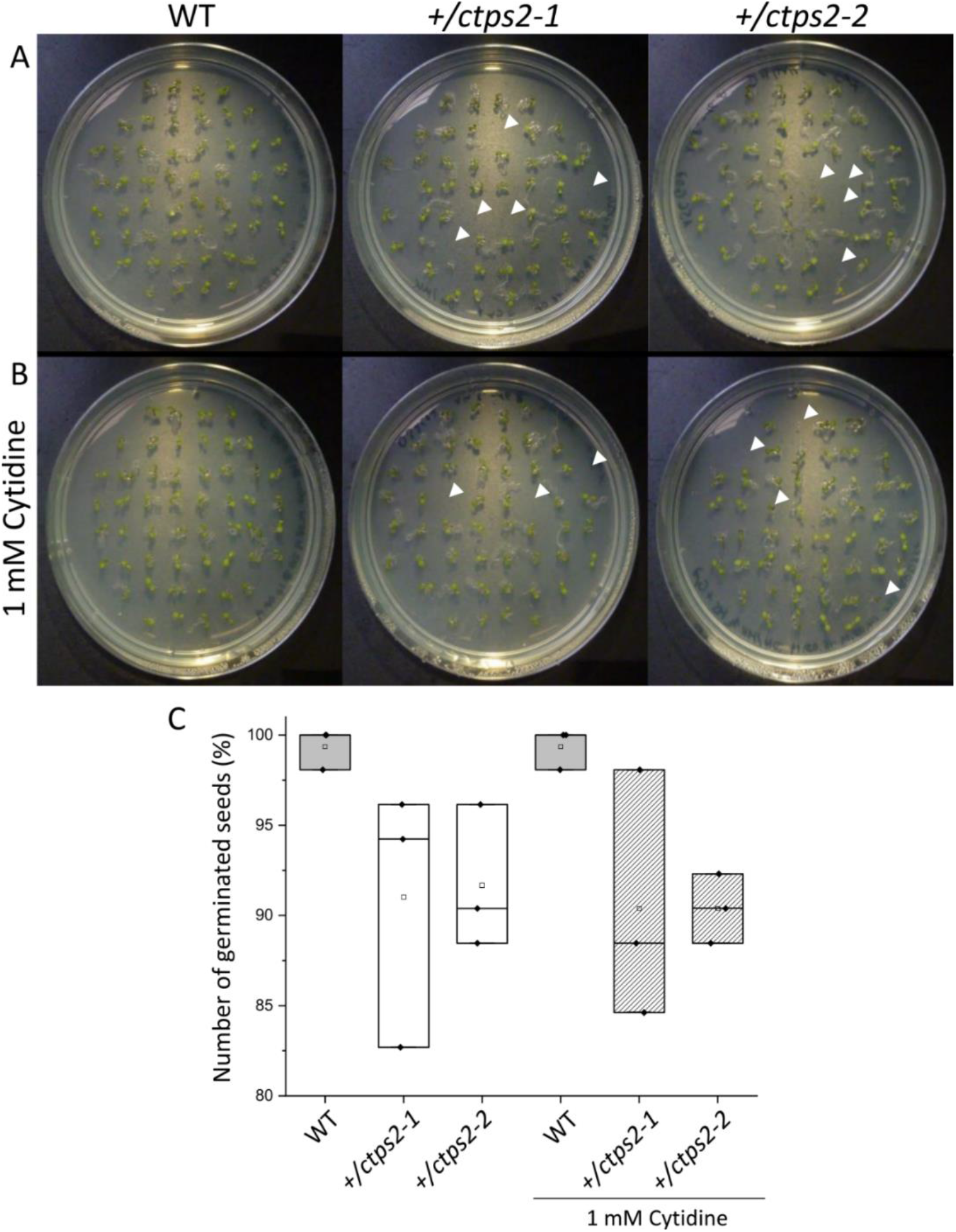
Germination of *+/ctps2* lines on ½ MS agar plates. WT and *+/ctps2* seeds were placed on ½ MS agar plates (A) or on plates supplemented with 1 mM cytidine (B). White arrowheads indicate non germinated seeds 5 days after transferring to growth chambers. Number of germinated seeds with and without application of 1 mM cytidine (C). A total of 52 seeds were plated for each genotype and the experiment was repeated three times.

### Siliques of *+/ctps2* plants contain nearly 25% aborted seeds

To investigate the seed development in detail, WT and *+/ctps2-1* seeds were sown on soil and single plants were separated to individual pots. Heterozygous T-DNA insertion in *+/ctps2-1* was checked by PCR. After the plants started reproductive growths, WT and *+/ctps2-1* siliques were harvested 8 days after fertilization (DAF). WT siliques contained healthy seeds with a deep green colored embryo inside, whereas *+/ctps2-1* siliques showed beneath healthy seeds, transparent embryo free as well as collapsed brown colored seeds (Figure 2A). To check whether supplementation with pyrimidines can rescue the phenotype, the blossoms of three *+/ctps2-1* plants was removed except one inflorescence with closed buds. These plants were irrigate with water containing 10 mM cytidine until siliques attained 8 DAF. Nevertheless, embryo free and collapsed seeds were still observable (Figure 2A), implicating that cytidine was not transported into the embryo or that the embryo is unable to recycle this compound. We assumed that the aborted seeds are homozygous for the T-DNA insertion and started to count seeds from *+/ctps2* plants. In both lines seeds were found, which were completely collapsed or had a nearly normal size, but were flat (Figure 2B). Both types were considered as aborted seeds. From a total of 2125 seeds in WT, 0.4% were aborted, whereas *+/ctps2-1* contained 22.3 % and *+/ctps2-2* 21.9 % aborted seeds in 2142 and 2475 seeds, respectively (Figure 2C).

**Figure 2.**
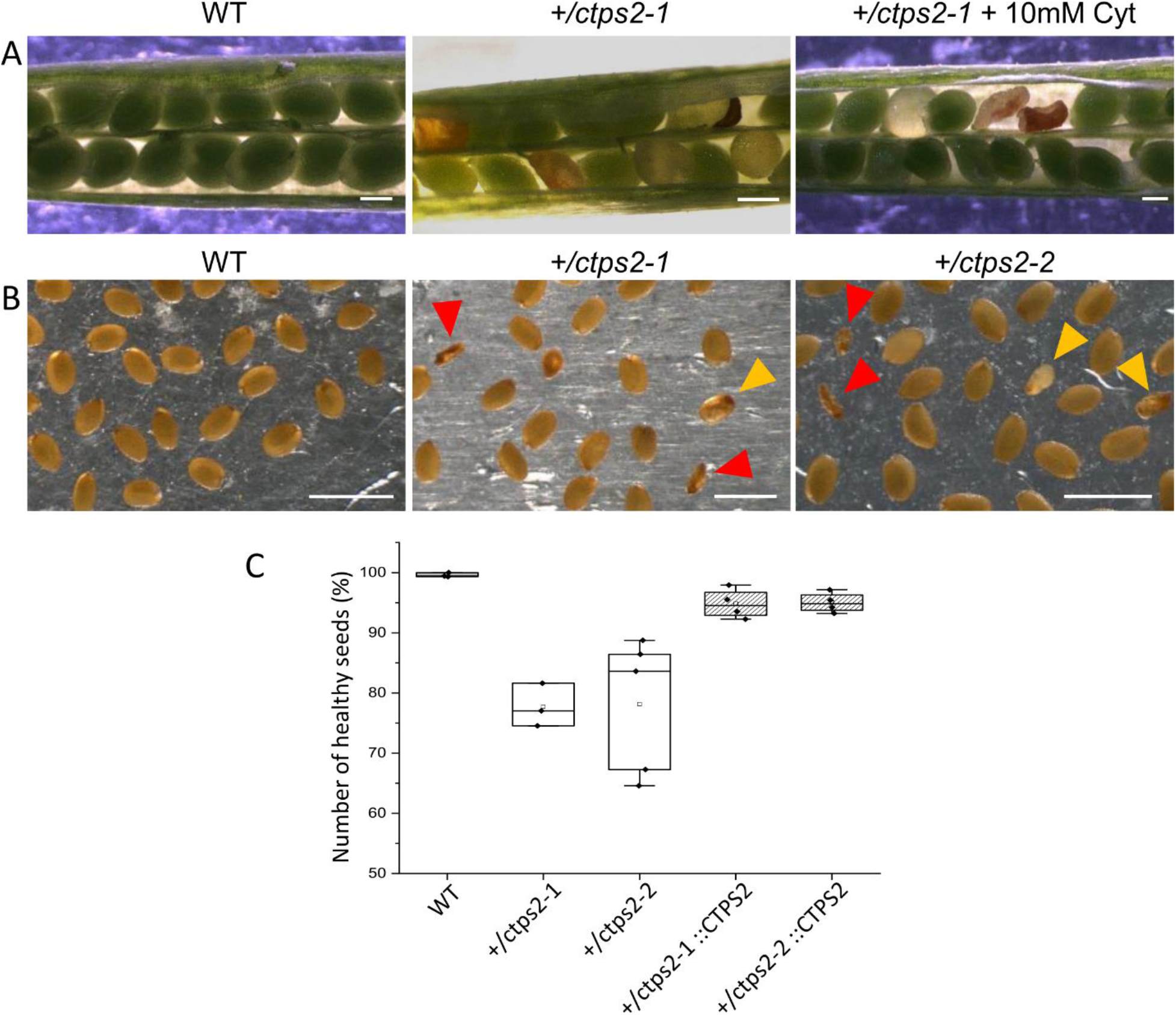
Identification of homozygous *ctps2* seeds. Siliques were opened 8-10 DAF from WT and *+/ctps2-1* plants. Irrigation of *+/ctps2-1* plants with 10 mM cytidine cannot rescue the aborted seed phenotype (A). 12 DAF siliques were harvested, desiccated at room temperature and seeds were collected using light microscopy. Red arrowheads point to collapsed seeds, while golden arrowheads indicate embryo free seeds with nearly normal size (B). Counting of seeds for WT, *+/ctps2* and complemented *+/ctps2* plants. Scale bar = 0.25 mm in (A) and 1 mm in (B). n= 3 for WT and *+ctps2-1* with a total of 38 and 40 siliques, respectively; n= 4 for *+/ctps2-1*::CTPS2 and *+/ctps2-2::CTPS2* with a total of 38 and 36 siliques, respectively; n= 5 for *+/ctps2-2* with 52 siliques in (C).

Integrating the full protein coding sequence of *CTPS2* under control of the full endogenous promoter (1864 bp upstream of ATG) into *+/ctps2* plants resulted in an average of only 5 % aborted seeds in both lines demonstrating, that the phenotype was at least partially complemented (Figure 2C). Attempts to rescue *+/ctps2* seeds phenotype by expression of other CTPS isoforms driven by UBQ promoter by us were not successful. Thus CTPS2 driven by the endogenous promoter seems to be crucial for proper embryo development.

### Reporter gene analysis reveals *CTPS2* promoter activity during embryo development

The visualization of *CTPS2* promoter activity used 1002 bp upstream of exon 1 and included a truncated promoter sequence compared to the complementation experiment. This nucleotide sequence was cloned in pBGWFS7.0 (Karimi et al., 2002) and fused to an eGFP and β-glucuronidase (GUS) tag after expression. Five positive transformants were identified and genomic integration of the construct was confirmed by PCR. Fertilized ovules at approximately 1.5 DAF showed strong *CTPS2* promoter activity in the peripheral and chalazal endosperm, but with strongest fluorescence signal in the peripheral endosperm (Figure 3). The *Arabidopsis* embryo develops to the heart stage 6 DAF and from this it develops quickly to the mature embryo until 8 DAF (Feng and Ma, 2017). Thus we focused on the promoter activity during rapid development rather than the globular stage. In the early heart stage we observed a punctate expression pattern, preferentially in embryos columella cells. Even in lateral view, rotated 90°, the late heart stage embryo showed the same expression pattern, but the fluorescence signal was specifically localized to only two or four root tip cells (Figure 3). When the embryo develops to the bent cotyledon stage the fluorescence signal remains strong in these columella cells. When the embryo reaches the mature stage *CTPS2* promoter activity was observed also in other tissues than the columella additionally. Herein promoter activity was observable in both cotyledons (Figure 3). Surprisingly the strong fluorescence signal in embryo columella cells seems to occur only in two cells, where the former connection between embryo and suspensor cells has been (Figure 4A). The early embryo and also the endosperm cells are connected to the maternal tissue via the suspensor to ensure nutrient and phytohormone supply by the mother plant (Bozhkov et al., 2005; Robert et al., 2018). Moreover, a maximum projection of the mature embryo revealed, strong *CTPS2* promoter activity in root epidermis cells and also in the cotyledons (Figure 4B). Especially in root epidermis cells it became obvious, that the fluorescence signal was not distributed homogenously, but seemed to be present in later atrichoblast or trichoblast cells (Figure 4B). These findings let us suggest, that *CTPS2* is spatiotemporal expressed in columella cells of the developing embryo. After 7 DAF the expression extends to further root and cotyledon tissues.

**Figure 3.**
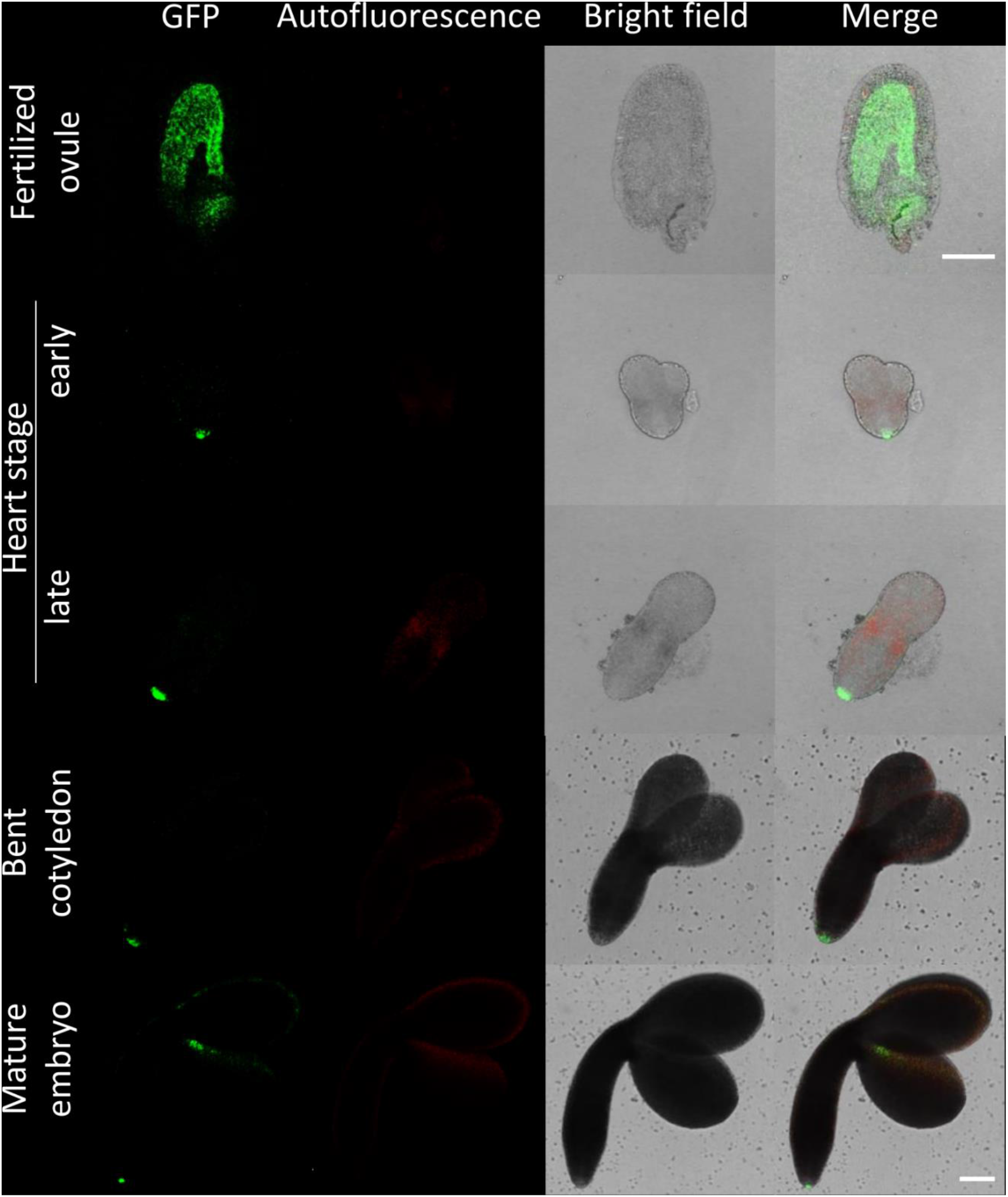
CTPS2 promoter activity in embryo cells. Siliques of soil grown plants were opened 2 DAF and between 6 and 8 DAF and seeds were harvested. Ovules and embryos were isolated from seeds for confocal laser scanning microscopy. Scale bar = 100μm.

**Figure 4.**
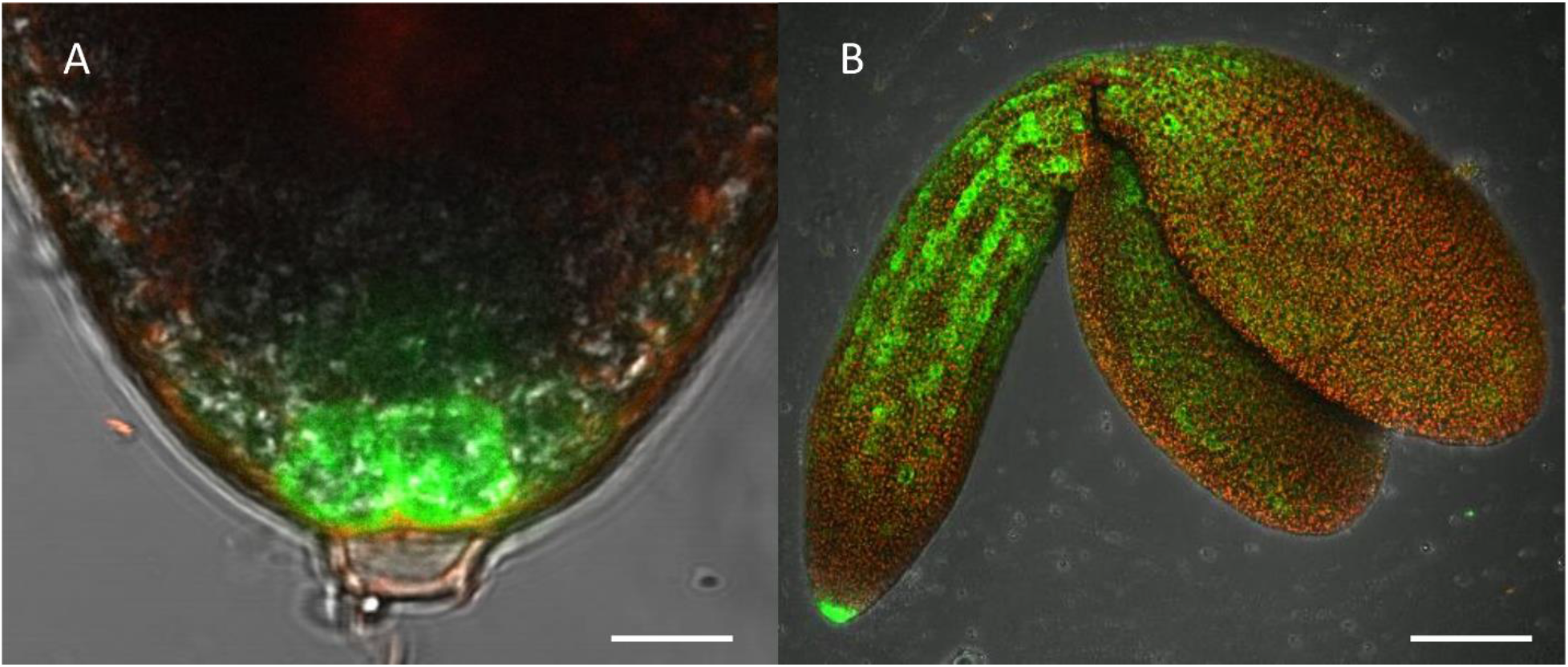
Highlight of CTPS2 promoter activity. Embryos were isolated from 8 DAF siliques for confocal laser scanning microscopy. Scale bar 10 μm in (A) and 100 μm in (B). Maximum projection of 24 pictures in (B).

### *CTPS2* promoter activity in seedling roots is restricted to columella cells and trichoblasts

We further investigated *CTPS2* promoter activity in the roots of young seedlings. One day after germination the fluorescence signal was very strong in root tip cells (Figure 5). In contrast to the images we took from developing embryos, roots were stained with the cell wall intercalating dye propidium iodide. Thereby, we found that *CTPS2* promoter activity is restricted to two central cells in the root tip (Figure 5). While the fluorescence signal intensity in central root tip cells decreased over time, it was completely absent from this cell type five days after germination (Figure 5). When the seedling becomes five days and older, the fluorescence signal shifts towards the elongation and later on towards the differentiation zone (Figure 5 and 6). In the differentiation zone *CTPS2* promoter activity was only found in trichoblast and developing root hair cells (Figure 6 and Figure S1). Since trichoblasts are able to develop into root hair cells, this finding is coherent. Mostly *CTPS2* promoter activity was restricted to young, developing root hair cells, pointing again to a strong spatiotemporal expression of CTPS2 (Figure 6 and Figure S1). By observing the primary root towards the shoot, a fluorescence signal was only detected in some trichoblasts, possibly soon developing into root hairs. Interestingly, we found strong *CTPS2* expression again in the root tip cells of lateral roots, emphasizing the observed strong spatiotemporal expression (Figure 6 and Figure S2).

**Figure 5.**
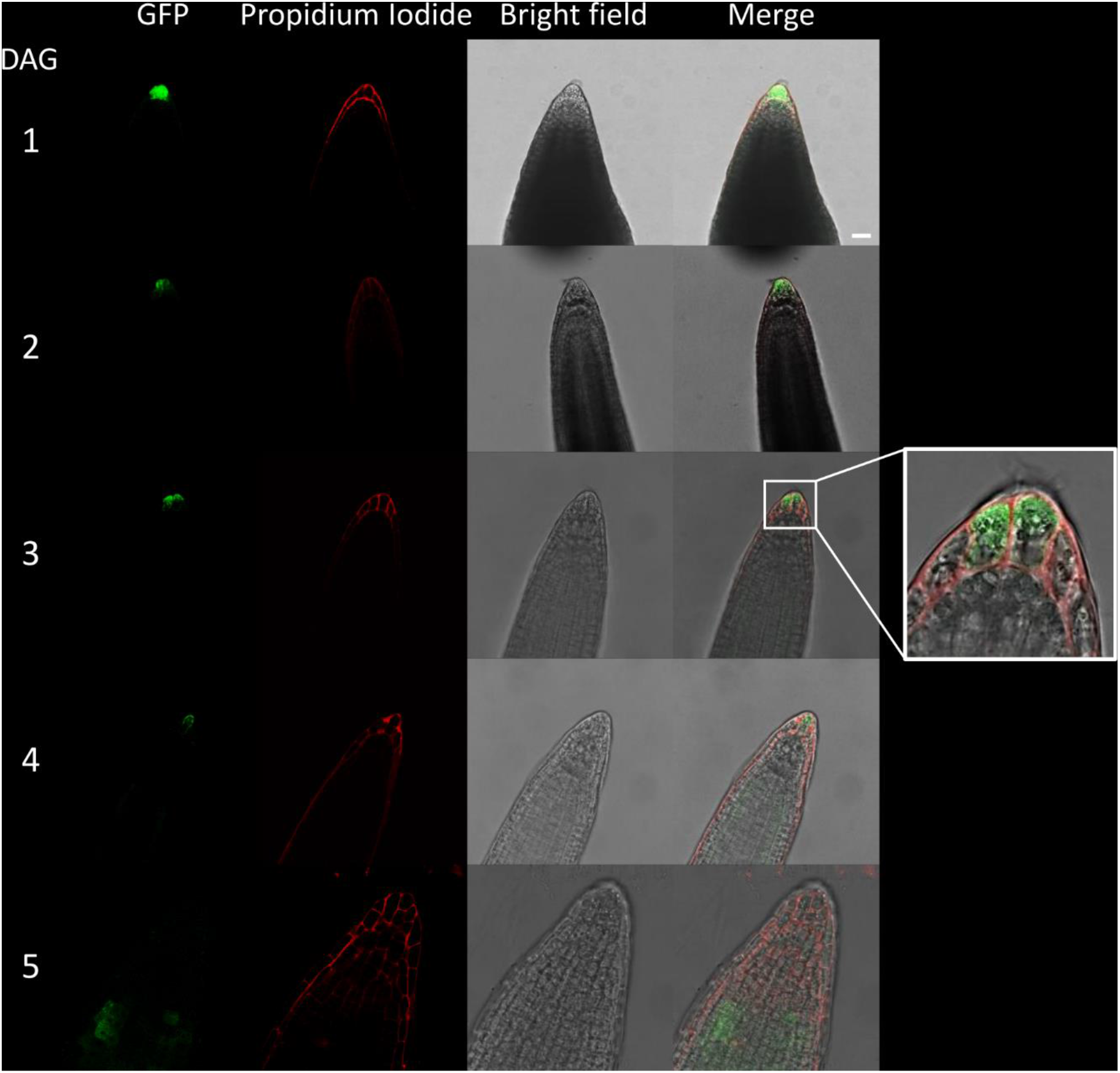
*CTPS2* promoter activity in young seedling roots. Seedlings were grown on ½ MS Agar plates and at the indicated time points transferred to slides with propidium iodide containing water for cell wall staining and subsequent confocal laser scanning microscopy with a Leica TCS SP5II microscope. Scale bar = 25 μm

**Figure 6.**
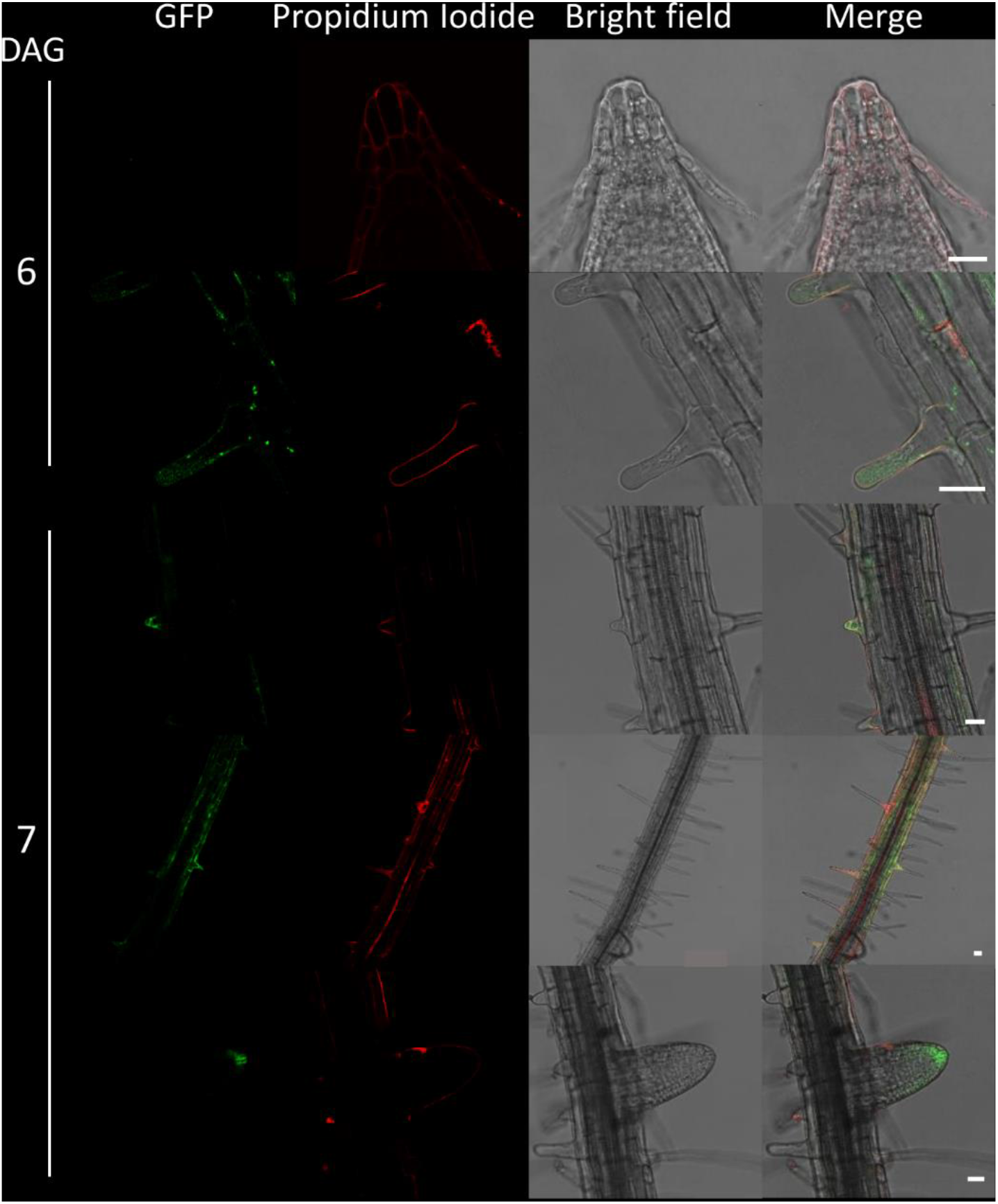
*CTPS2* promoter activity in roots. Growth and microscopy conditions as in figure 5. Scale bar = 25 μm

### Auxin treatment increases *CTPS2* promoter activity

Since the *CTPS2* promoter activity in columella cells revealed a similar fluorescence signal, compared to the established auxin reporter DR5, we aligned the nucleotide sequence of the auxin response element (TGTCTC) to the *CTPS2* promoter sequence used in this study (Ulmasov et al., 1995, Ulmasov et al., 1997; Feng and Ma 2017). Within the alignment we found a repeat of the auxin response element (AuxRE) in the *CTPS2* promoter at position 652-665 upstream of exon 1 interrupted by the two bases CG (Figure S3). This finding gives a hint, that *CTPS2* may be auxin regulated. To support this finding, the two *CTPS2* reporter lines # 10 and 15 were grown on ½ MS agar plates for 7 days and transferred to ½ MS agar plates supplemented with 250 nM NAA or 250 nM DMSO. After 20 h growth, confocal laser scanning microscopy with subsequent fluorescence intensity analysis, using ImageJ (version 1.51j8), revealed indeed a stronger GFP signal in trichoblasts of NAA treated plants (Figure S4). Under control conditions the mean grey value per cell of line # 10 and 15 were 184.7 and 159.4, respectively. NAA treatment increased the mean grey value per cell in both lines significantly, resulting in 216.8 for line # 10 and 220.8 for line # 15 (Figure 7). Although the increase in fluorescence intensity was most prominent in trichoblast cells in both lines, NAA treatment induced fluorescence also in cortex cells. However, no significant upregulation of fluorescence intensity was found for cells within the root cortex (data not shown).

**Figure 7.**
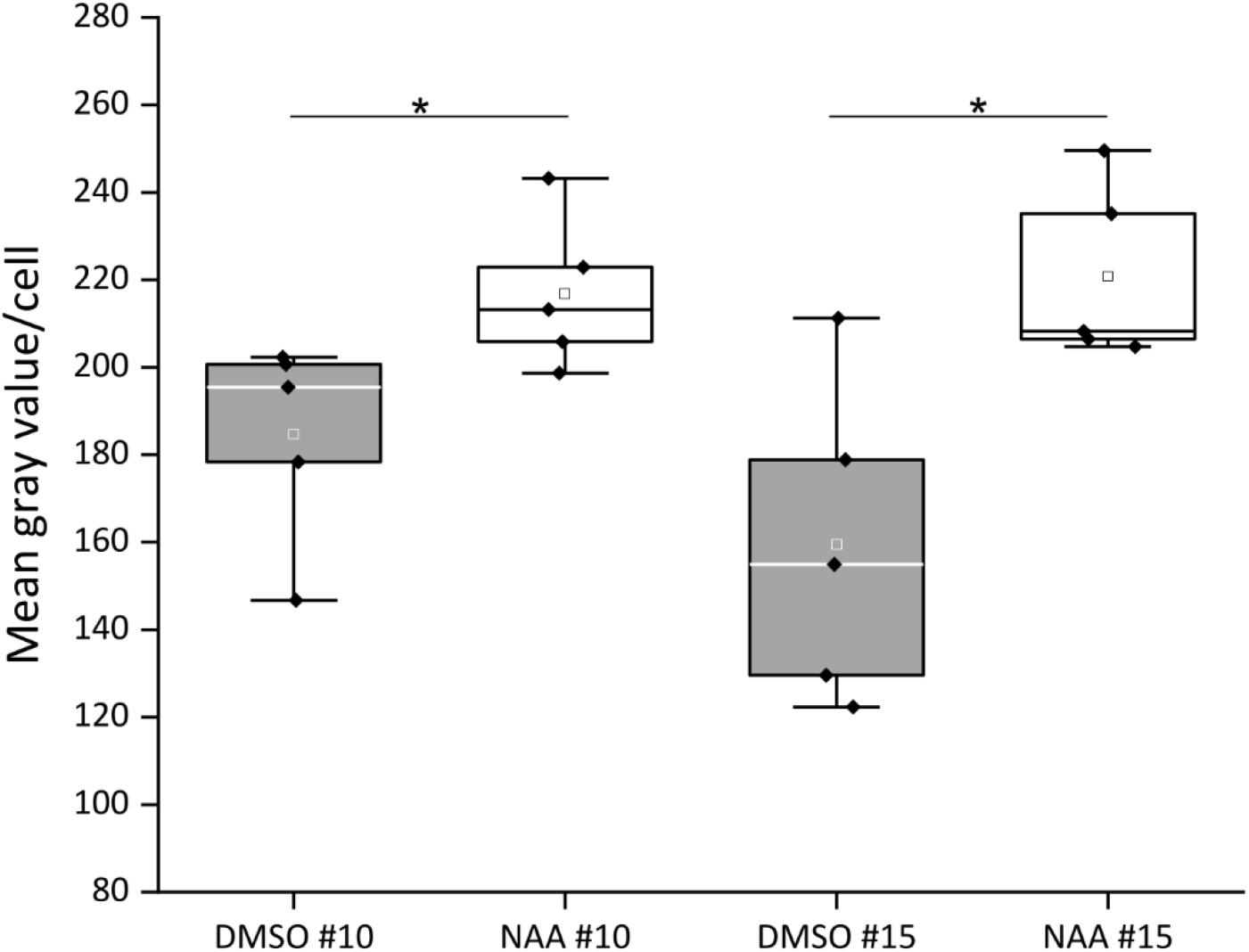
CTPS2 promoter fluorescence intensity after NAA application. Five day old seedlings, grown on ½ MS agar plates were transferred to 250 nM NAA or control (DMSO) plates for 20h. The fluorescence intensity of 5 biological replicates with10 cells per replicate are represented in box plots. White and grey rectangle represent mean value in DMSO and NAA treated plants, respectively. Median is shown as white or black line in DMSO or NAA box plot, respectively. Statistical analysis used Student’s t-Test with p <0.05.

## Discussion

During plant development, especially young tissues require enormous amounts of nucleotides for cell division and growth. Therefore, it is assumed, that *de novo* synthesis rates in seedlings and growing tissues are generally high (Zrenner et al., 2006). This hypothesis was supported by the finding of strong ATCase promoter activity, the enzyme which commits the rate limiting step in pyrimidine *de novo* synthesis, in Arabidopsis seedlings (Bassett et al., 2003; Chen and Slocum, 2008DR5). A decrease in nucleotide *de novo* synthesis, caused by lower UMP synthase expression, increases pyrimidine salvage activity in growing potato tubers (Geigenberger et al., 2005). In contrast to that, differentiated cells maintain their nucleotide pools via salvage and compensate catabolic processes by minor *de novo* synthesis activity (Zrenner et al., 2006). Due to that, the external supplied nucleoside cytidine, which can be imported in plants by ENT3 (Traub et al., 2007) and can by phosphorylated to CMP by uridine/cytidine kinase (Ohler et al., 2019) should allow for a compensation of lost CTP synthase activity in *ctps2* seedlings. However, the application of 1 mM cytidine did not overcome the germination phenotype of *ctps2* seeds (Figure 1). Together with the finding, that siliques from *+/ctps2* plants contain nearly 25 % aborted seeds, which also cannot be rescued by irrigation with 10 mM cytidine containing water, but by complementation of *+/ctps2* plants with CTPS2 under control of the endogenous promoter, we conclude that the crucial function of CTPS2 lays in the embryo development (Figure 2) and corresponding salvage activity or import competence for cytidine is lacking in this tissue.

It is known from work on drosophila, that egg chamber development goes along with a high demand for nucleotides to allow for an increased rRNA synthesis () Surprisingly, we found visible *CTPS2* promoter activity only in the tip of columella cells in embryos of the heart and later stages. During embryo development *CTPS2* promoter activity was observed in other embryonic tissues after 7 DAF (Figure 3 and 4). In contrast to that, we noticed a relatively strong CTPS2 promoter activity in the peripheral endosperm of ovules 1.5 DAF (Figure 3, upper panel). Together with the finding of collapsed and embryo free seeds in 8-10 DAF siliques of *+/ctps2-1* plants it is likely, that a knockout of *CTPS2* causes embryonic growth arrest at early embryo development. Andriotes and Smith (2019) found that impaired purine synthesis causes growth arrest and embryo abortion at the globular stage. This was also found for genes in the oxidative pentose phosphate pathway, providing precursors for phosphoribosyl-pyrophosphate (a co-substrate in nucleotide de novo synthesis) synthesis and UMP synthase. Thus we think, that *CTPS2* is an *EMB* gene, causing growth arrest at a similar developmental stage. Moreover, the knockout of enzymes in the pyrimidine *de novo* synthesis, which are encoded by a single gene is embryo lethal (Chen and Slocum, 2008; Schröder et al., 2005). Since Arabidopsis possesses five CTP synthase genes, a spatiotemporal regulation of these is likely and would explain why *CTPS2* knock-outs cannot be complemented by other CTP synthase isoforms. Indeed, no other Arabidopsis CTP synthase harbors a repeat of the AuxRE in its promoter, except *CTPS2*. However, there is still the question, why *CTPS2* promoter activity is still restricted to the columella tip cells in the developing embryo? Stadler et al., (2005) established a mobile and membrane bound GFP approach under control of the *At*SUC3 promoter, which allowed monitoring of cell-to-cell movement. *At*SUC3 promoter activity was described in several sink tissues as well as the suspensor (Meyer et al., 2004). Nevertheless, Stadler et al., (2005) observed GFP fluorescence signals in embryos at the globular stage, concluding that the suspensor and early embryo are building a symplastic system. After the disconnection of suspensor and embryo (Bozhkov et al., 2005) and progression in embryo development, GFP fluorescence becomes more and more restricted to individual cell types, e.g. epidermis cells (Stadler et al., 2005). This implicates, that symplastic movement of macromolecules is no longer facilitated in the older embryo, but a tissue specific connectivity by plasmodesmata must still remain (Stadler et al., 2005). It is accepted, that molecules of a mass of up to 800 Da are able to diffuse freely via symplastic transport (Karmann et al., 2018; Lalonde et al., 2004). Since CTP has a mass of 483 Da it is conceivable, that CTPS2 produces CTP in embryonic columella cells, which is symplastically distributed between all embryonic cells. The reduction of plasmodesmata during further development triggers CTPS2 activity in other cell types, resulting in GFP fluorescence in epidermis cells and cotyledons (Figure 4B).

The *CTPS2* expression pattern was similar to that of the auxin reporter DR5::GFP (Feng and Ma, 2017) during embryogenesis and an AuxRE repeat was located in the *CTPS2* promoter region (Figure S4). Maternal tissues supply the embryo with auxin until the globular stage, when the embryo is able to produce auxin by itself (Robert et al., 2018). With the fertilization of the ovum, suspensor cells express PIN7 for a directed auxin transport towards the embryo and the auxin concentration is high at the embryos basal end (Friml et al., 2003). When the embryo starts independent auxin production in its apical cells, PIN1 expression is activated, resulting in an apical-basal auxin gradient, accumulating auxin in the basal end of the embryo and in upper suspensor cells (Friml et al., 2003; Robert et al., 2013; Robert et al., 2018; Vieten et al., 2005). Growth is a high demand time for nucleotides, thus, coupling of CTPS expression to the growth hormone auxin apparently makes sense. Symplastic coupling of embryonic cells could then allow for a distribution of nucleotides (CTP) within the embryo. At the same time, such system would possibly allow for better regulation compared to a salvage of phloem derived cytidine, which according to our results does not take place.

In the root, DR5::GFP is expressed in the primary- and lateral root tip, especially in columella cells, similar to the observations made in this study (Figure 5 and 6 (Wabnik et al., 2011; Raya-González et al., 2018)). DR5::GFP is a very strong sensor, which uses 9 inverted repeats of a 11bp sequence, containing the TGTCTC sequence (Ottenschläger et al., 2003; Ulmasov et al., 1997). Therefore it is conclusive, that GFP fluorescence signals observed under control of the *CTPS2* promoter are not as intensive as DR5::GFP signals, but can be intensified by auxin application (Figure 7, Figure S4). However, the columella is a highly dynamic tissue, which probably demands an extensive amount of CTP. Beneath its use as building block for RNA and DNA, in yeast CTP is an essential precursor in membrane-phospholipid biosynthesis. Although CTP inhibits CTPS enzyme activity, in yeast it was shown, that the phosphorylation of CTPS stimulates its activity and increases endogenous CTP concentrations (Chang and Carman, 2008). Since the posttranscriptional regulation mechanisms and biochemical properties of plant CTPS are not completely understood yet (Daumann et al., 2018), it is possible that CTPS from Arabidopsis could be regulated similarly to yeast CTPS.

Root hair formation is initiated by auxin. The phytohormone stimulates AuxRE and thereby induces trichoblast specific kinases, which initiate trichoblast elongation (Vissenberg et al., 2020). The major constituent of plant membranes is phosphatidylcholine and its *de novo* synthesis requires CDP-choline, delivered from the Kennedy pathway. One of the initial steps is the transfer of the cytidyl moiety from CTP to phosphocholine (Caldo et al., 2019; Kennedy, 1958). The transition from a trichoblast cell into a root hair, which is in the end still one cell, requires enormous amounts of membrane phospholipids and thus CTP. Together with our finding of high *CTPS2* promoter activity in trichoblasts and root hair cells (Figure 6, Figure S1), *CTPS2* transcript was found to be upregulated in a RNA-seq analysis on Arabidopsis root hair cells (Huang et al., 2017).

Taken together, we conclude from our data, that *CTPS2* is an *EMB* gene, regulated by auxin and its knockout is embryo lethal. Moreover, *CTPS2* is auxin-dependent expressed in root columella cells and trichoblasts and may contribute to their development.

## Author Contributions

All authors contributed to the research design and wrote the manuscript. D.H. generated complementation and promoter-reporter-gene lines and analyzed seed development and germination. D.H. and D.S. performed confocal microscopy. D.S. advised Auxin treatment experiments and quantification of fluorescence intensity.

