## Supplemental Table 1 and Supplemental Figures 1-4 for "CTP-Synthase 2 from *Arabidopsis thaliana* is required for complete embryo development"

**Hickl et al.**

**Supplemental Tables**

Table S1: Primers used in this study

| Designation | s-sequence (5’-3’) | as-sequence (5’-3’) | Description |
|---|---|---|---|
| CTPS2_Promoter | attgttcatcaccactcttctctc | aatcgtttttgtctctgcttcttc | gDNA amplification |
| CTPS2_Full-length | aatttgaaccttgggctggttaca | gtggtgaagcccgtttccattg | gDNA amplification |
| CTPS2_Promoter | ggggacaagtttgtacaaaaaagcaggcttaattgttcatcaccactctt | ggggaccactttgtacaagaaagctgggtaaatcgtttttgtctctgctt | att-site attachment |
| CTPS2_Full-length | ggggacaagtttgtacaaaaaagcaggcttaaatttgaaccttgggctgg | ggggaccactttgtacaagaaagctgggtagtggtgaagcccgtttccat | att-site attachment |
| *+/ctps2-1* | gtaagtcttccttgaccctaagcg | | Left border |
| *+/ctps2-1* | aaaagctcatgaagcgatcctaag | | Right border |
| *+/ctps2-2* | gctcatatctccgtcgttacaaac | | Left border |
| *+/ctps2-2* | ccaggtatttgtactacctcctca | | Right border |
| GK_o8474 | ataataacgctgcggacatctacatttt | | LB-primer |
| CTPS2_700 | atgatatcttgttgtaatcctagt | | Sequencing |
| CTPS2_1400 | atgcaaacttccacaacacaaaca | | Sequencing |
| CTPS2_2100 | gtagctatgagatcctttttcaca | | Sequencing |
| CTPS2_2800 | acaggtgatattgaatctatg | | Sequencing |
| CTPS2_3500 | gttcctctgcttttaagggt | | Sequencing |
| CTPS2_4200 | catagagtgggttgcagcta | | Sequencing |
| CTPS2_4900 | aaatctatataacttctaatctct | | Sequencing |
| CTPS2_5600 | ctcgagaaaggaacaattttactg | | Sequencing |
| Ef1α | gagaccaccaagtactactgcac | gttggtcccttgtaccagtcaag | Quality control |

### Supplemental Figures

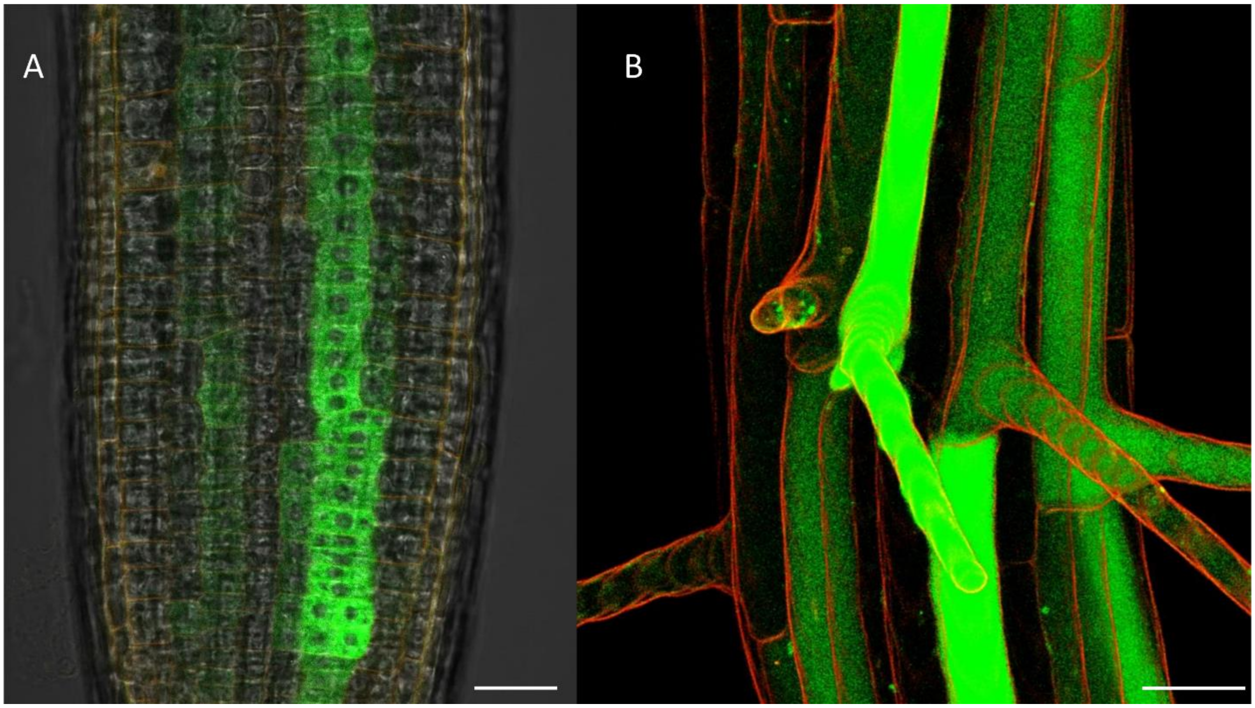


**Figure S1. CTPS2 promoter activity is strong in trichoblasts and root hair cells.** Plants were grown on ½ MS agar plates, transferred to slides and cell walls were stained with propidium iodide. Confocal laser scanning microscopy was conducted with a Zeiss LSM880 AxioObserver SP7. Scale bar 25 µm. (B) 39 pictures were used as maximum projection with 4 µm frame for each picture.

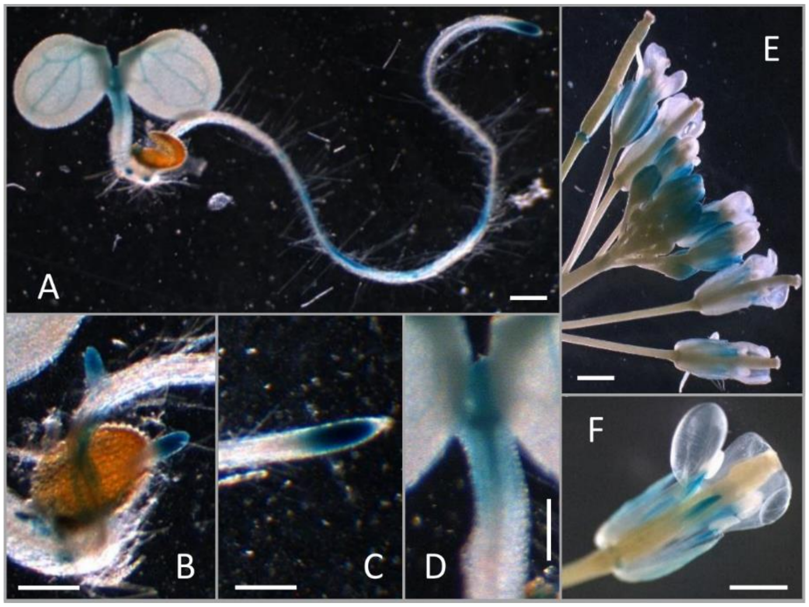


**Figure S2. CTPS2 promoter activity in seedlings and in reproductive tissues.** (A-D) Five day old seedlings grown on ½ MS agar plates. (A) Seedling with promoter activity in the primary root, shoot apical meristem and vasculature tissue. (B, C) Root tips of primary (C) and secondary roots (B) have strong CTPS2 promoter activity. (D) CTPS2 promoter activity is moderate in the shoot apical meristem. (E, F) During reproductive growth phase *CTPS2* promoter activity occurs in filaments of the flower. Scale bar = 1 mm

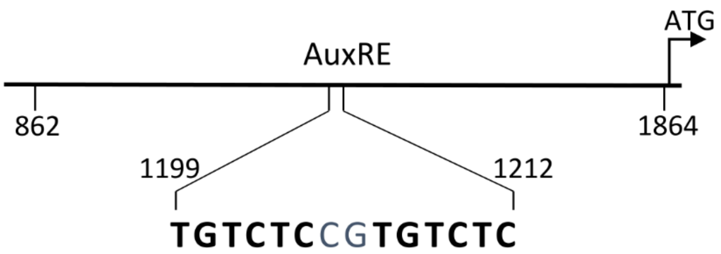


**Figure S3. Auxin response element found in the promoter region of *CTPS2*.** The sequence was included in all promoter activity studies conducted in this work.

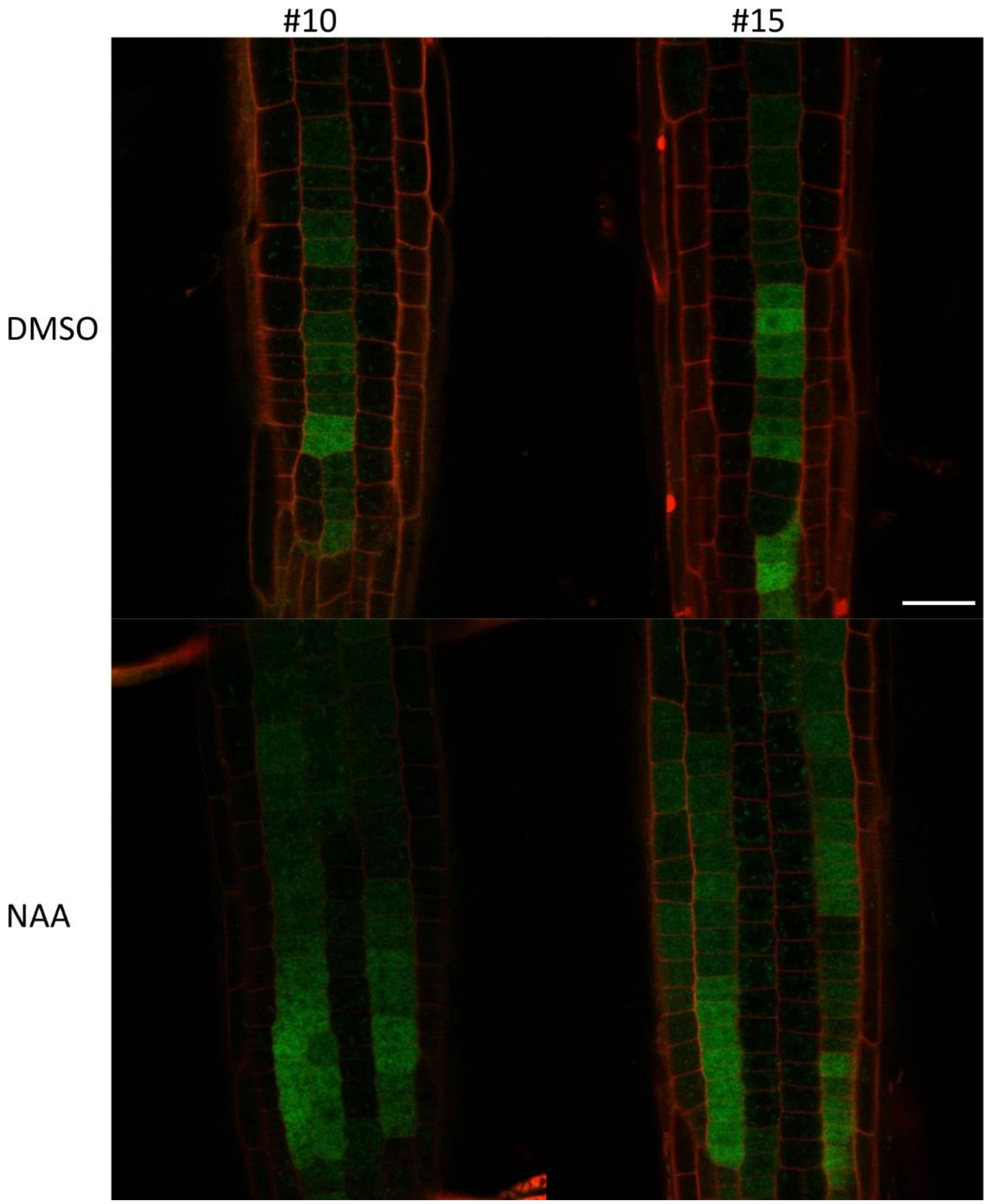


**Figure S4. NAA treatment increases CTPS2 promoter activity.** Seedlings of the two independent CTPS2 reporter lines # 10 and 15 were grown on ½ MS agar plates for 5 days and transferred to plates containing DMSO as control or 250 nM NAA for 20 h. NAA treatment increased the GFP signal specifically in trichoblast cells. Propidium iodide was used to stain cell walls. Scale bar = 25 µmm.
